# NR5A2 connects genome activation to the first lineage segregation in early mouse development

**DOI:** 10.1101/2022.11.25.518012

**Authors:** Fangnong Lai, Lijia Li, Xiaoyu Hu, Bofeng Liu, Ziqi Zhu, Ling Liu, Qiang Fan, Huabin Tian, Kai Xu, Xukun Lu, Qing Li, Feng Kong, Lijuan Wang, Zili Lin, Hongyu Deng, Jinsong Li, Wei Xie

## Abstract

After fertilization, zygotic genome activation (ZGA) marks the beginning of the embryonic program for a totipotent embryo, which further gives rise to the pluripotent embryonic lineages and extraembryonic trophectoderm after the first lineage commitment. While much has been learned about pluripotency regulation, how ZGA is connected to the pluripotency commitment in early embryos remains elusive. Here, we investigated the role of nuclear receptor ^1^ family transcription factors (TFs) in mouse pre-implantation embryos, whose motifs are highly enriched in accessible chromatin at the 2-cell (2C) to 8-cell (8C) stages after ZGA. We found NR5A2, an NR TF highly induced upon ZGA, is required for early development, as both the knockdown and knockout of *Nr5a2* from 1C embryos led to morula arrest. While the zygotic genome was largely activated at the 2C stage, 4-8C-specific gene activation (mid-preimplantation activation) was substantially impaired. Genome-wide chromatin binding and RNA-seq analyses showed NR5A2 preferentially regulates its binding targets including a subset of key pluripotency genes (i.e., *Nanog, Pou5f1*, and *Tdgf1*). Finally, NR5A2-occupied sites at the 2C and 8C stages predominantly reside in accessible B1 elements where its motif is embedded. Taken together, these data identify NR5A2 as a key regulator that connects ZGA to the first lineage segregation in early mouse development.

## Introduction

Following fertilization, the global transcription of the embryo is initially quiescent.^2^ After a few cell cycles, thousands of zygotic transcripts are produced, known as zygotic genome activation.^3^ ZGA can be divided into two waves: the minor ZGA and major ZGA, which occur at the middle 1-cell (1C) stage and late 2-cell (2C) stage in mice, respectively.^4^ After ZGA, the embryos give rise to two lineages: inner cell mass (ICM), which includes the founder of pluripotent cells, and trophectoderm (TE), which develops to the extraembryonic layer.^5^ ICM subsequently gives rise to *Nanog*-expressing pluripotent epiblast and *Gata6*-expressing primitive endoderm at E4.5.^6^ The epiblast cells are the origin of all future embryonic lineages.^7^ The naïve pluripotency can be captured *in vitro* through the derivation of ESCs from ICM. The core pluripotency TFs OCT4, SOX2 and NANOG cooperatively regulate the intrinsic pluripotency transcription network.^6, 8-10^ Besides, ESCs also express “ancillary” pluripotency regulators, such as ESRRB, KLF4, SALL4, and TBX3, which can enhance the pluripotency network.^5^ Despite remarkable progress in understanding the pluripotency regulation, how ZGA is connected to the pluripotency establishment remains elusive to date.

Recently, with the advance of ultra-sensitive chromatin analysis approaches, dramatic reprogramming of chromatin landscapes has been revealed in early mammalian embryos.^3, 11^ These chromatin maps also identified putative regulatory elements that exhibit highly dynamic activities in early embryogenesis, which may recruit master TFs to regulate the transcription network. However, the identities of these TFs and how their interactions with *cis*-regulatory elements govern transcription circuitry remain poorly understood. Notably, TFs can be inferred from motifs embedded in regulatory elements identified by transposase accessible chromatin with high-throughput sequencing (ATAC-seq) or DNase I hypersensitive site sequencing (DNase-seq).^1, 12^ Interestingly, the motifs of nuclear receptor ^1^ family factors such as NR5A2 and RARG are overrepresented in the accessible putative regulatory elements in mouse 2-8C embryos.^13^ NRs are structurally conserved, ligand-dependent TFs, with diverse functions in cell proliferation, metabolism, stem cell pluripotency, and homeostasis.^14^ For example, RARG was shown to participate in the 2C-like-cell program,^15^ although its deficiency alone was compatible with mouse early development likely due to redundancy from the RAR family.^16, 17^ NR5A2 was suggested to bind phosphatidyl inositol, although murine NR5A2 appears to have lost the ligand binding ability and becomes ligand-independent during evolution.^18^ NR5A2 is also closely linked to pluripotency regulation. *Nr5a2* is essential for early embryogenesis and regulates the expression of *Oct4* in epiblast.^19-21^ In mouse embryonic stem cells (mESCs), NR5A2 is regulated by canonical Wnt/b-Catenin signaling and in turn directly controls pluripotency genes *Oct4, Nanog*, and *Tbx3*.^22^ Furthermore, NR5A2 can induce epiblast stem cell (EpiSC) into ground-state pluripotency,^23^ and replace OCT4 in iPSC reprogramming.^20^ ESRRB and NR5A2 cooperatively play essential roles as auxiliary factors of OCT4 and SOX2 for the core-pluripotent network in mESCs.^24^ These studies underscore potential roles of NR TFs in early development and pluripotency regulation. However, how these TFs interact with chromatin and govern transcription circuitry *in vivo* remains poorly understood.

Here, we screened for possible NR TFs that may function in early embryos after ZGA and identified NR5A2 as a critical regulator for mouse early development. Knockdown (KD) or knockout (KO) of *Nr5a2* impaired the activation of 4-8C genes and led to embryonic arrest around the morula stage. Using CUT&RUN, we profiled NR5A2 chromatin occupancy at the 2C and 8C stages, and revealed that NR5A2 directly activated many genes including a set of key pluripotency genes. Furthermore, we found NR5A2 preferentially bound B1 elements near major ZGA genes. These data suggest NR5A2 plays a critical role in regulating early embryo development by connecting ZGA to the first lineage specification program.

## Results

### Nuclear receptor TFs regulate the mouse early development

To identify putative key transcription factors functioning between ZGA and lineage segregation in mouse early development, we firstly investigated the regulatory elements from our previous ATAC-seq data,^13, 25^ which showed strong enrichment of motifs of NR TFs, including NR5A2 and RARG, from the 2C to 8C stage (Fig. 1a). Consistently, these two TFs were highly activated upon ZGA but their expression declined after the 8C stage (Fig. 1b). As *Nr5a2* showed a weak level of transcript in oocytes, we confirmed that NR5A2 protein was undetectable in germinal vesicle (GV) oocytes but was strongly induced at the 2C stage using immunofluorescence (Fig. 1c). Given NR family members often share similar motif sequences,^26^ we asked if the NR family TFs may play redundant roles during early development. RNA-seq analysis showed that two additional NR family TFs *Nr1h3* and *Nr2c2* were also highly expressed from 2C to 8C, and exhibited similar binding motifs (Fig. 1b). Therefore, we first performed quadruple knockdown (4KD) against these four NR factors by injecting combined siRNAs into zygotes (Fig. S1a). Remarkably, the 4KD embryos developed to 8C at a lower rate and barely developed to blastocysts, with most embryos arrested at the morula stage (Fig. S1b). RNA-seq confirmed the knockdown efficiency of these NR TFs at the 4C and 8C stages (Fig. S1c). Compared with the control group, 4KD led to 103 down-regulated genes and 66 up-regulated at the 4C stage (fold change >2, P-value <0.05) (Fig. S1d and Methods). At the 8C stage, the down-regulated and up-regulated gene numbers increased to 281 and 173, respectively (Fig. S1d). In particular, 4C-activated genes and 8C-activated genes (Methods) were substantially down-regulated upon the KD of the 4 NR TFs (Fig. S1e). Taken together, these data indicate that NR family TFs play important roles in early development and transcription regulation at the 4-8C stage.

**Fig. 1.**
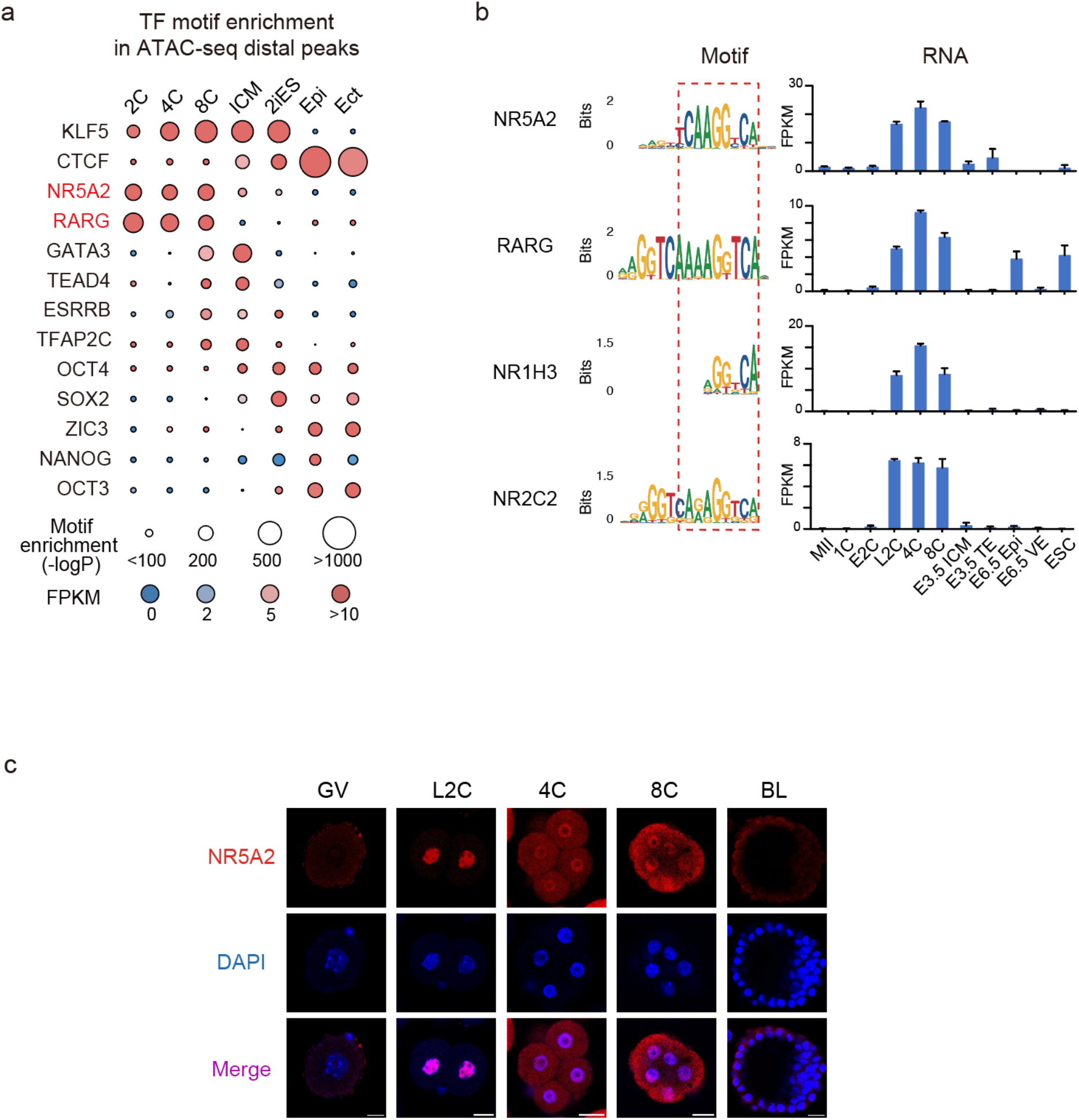
Nuclear receptor TFs in mouse early development. **a**, TF motifs identified from distal ATAC-seq peaks^13, 25^ in mouse early embryos. Sizes of circles indicate -Log P value. Expression levels of TFs are color coded. 2iES, 2i mESC; Epi, E6.5 Epiblast; Ect, E7.5 ectoderm. **b**, Sequence logos of the binding motifs of NR5A2, RARG, NR1H3, and NR2C2 (left), and RNA expression (FPKM) of *Nr5a2, Rarg, Nr1h3*, and *Nr2c2* (right). VE, visceral endoderm. PS, primitive streak; Mes, mesoderm; End, endoderm. The error bars denote the standard deviations of two biological replicates of RNA-seq. **c**, Immunofluorescence of NR5A2 (red) and DAPI (blue) in mouse germinal vesicle (GV) oocyte, L2C, 4C, 8C embryos and blastocyst. Scale bar: 20 μm.

### Knockdown or knockout of *Nr5a2* led to developmental arrest at the morula stage

To ask which NR TF plays a more critical role in early development, we removed each one of the four siRNAs in the combined siRNAs and evaluated the developmental rates at E4.5 (blastocyst) (Fig. S2a, b). Remarkably, when knocking down these 4 NR TFs, leaving out the siRNA targeting *Nr5a2*, but not *Nr1h3* or *Nr2c2*, substantially “rescued” the morula arrest phenotype of the 4KD embryos (Fig. S2c). Leaving out *Rarg* siRNA had a moderate “rescue” effect (Fig. S2c). Consistently, leaving out *Nr5a2* siRNA also restored normal 4C and 8C transcriptional programs (Fig. S2d), indicating NR5A2 is the main TF responsible for the 4KD phenotype. To further confirm this result, we performed single KD for each of the 4 NR TFs (Fig. 2a). Indeed, only knocking down *Nr5a2* resulted in a severe arrest at the morula stage (Fig. 2b-c). Knocking down *Rarg* led to a milder developmental defect (Fig. 2b-c). To rule out the possible knockdown off-target effects, we attempted to mutate *Nr5a2* via enhanced base editing (EBE) in embryos (Fig. S3a),^27^ which achieved high KO efficiency (96%) (Fig. 2d, top left, Fig. S3b). The depletion of NR5A2 protein was confirmed by immunofluorescence (Fig. 2e). Consistently, these mutant embryos arrested at the morula stage, thus phenocopying the knockdown of *Nr5a2* (Fig. 2d, top right and bottom). Finally, RNA-seq analysis showed knocking down *Nr5a2* led to the strongest transcription changes among the 4 NR TFs, with 976 down-regulated and 806 up-regulated genes at the 8C stage (Fig. S3c). Furthermore, among the 4 KD groups, only *Nr5a2* KD recapitulated the failed activation of 4-8C genes (Fig. 2f), with similar genes affected as in 4KD embryos (Fig. S3d). We noticed that after KD of *Nr5a2* (Fig. 2g, top left), the embryos could develop to 2C normally (Fig. 2g, top right and bottom) with the zygotic genome clearly activated (Fig. 2h, Fig. S3e). We did not observe an evident global activation defect as the numbers of ZGA genes that were upregulated (n=155) and downregulated (n=138) were relatively small and comparable (Fig. 2i). Taken together, these results suggest that NR5A2 is the dominant factor among the four NR TFs activated upon ZGA in regulating early development.

**Fig. 2.**
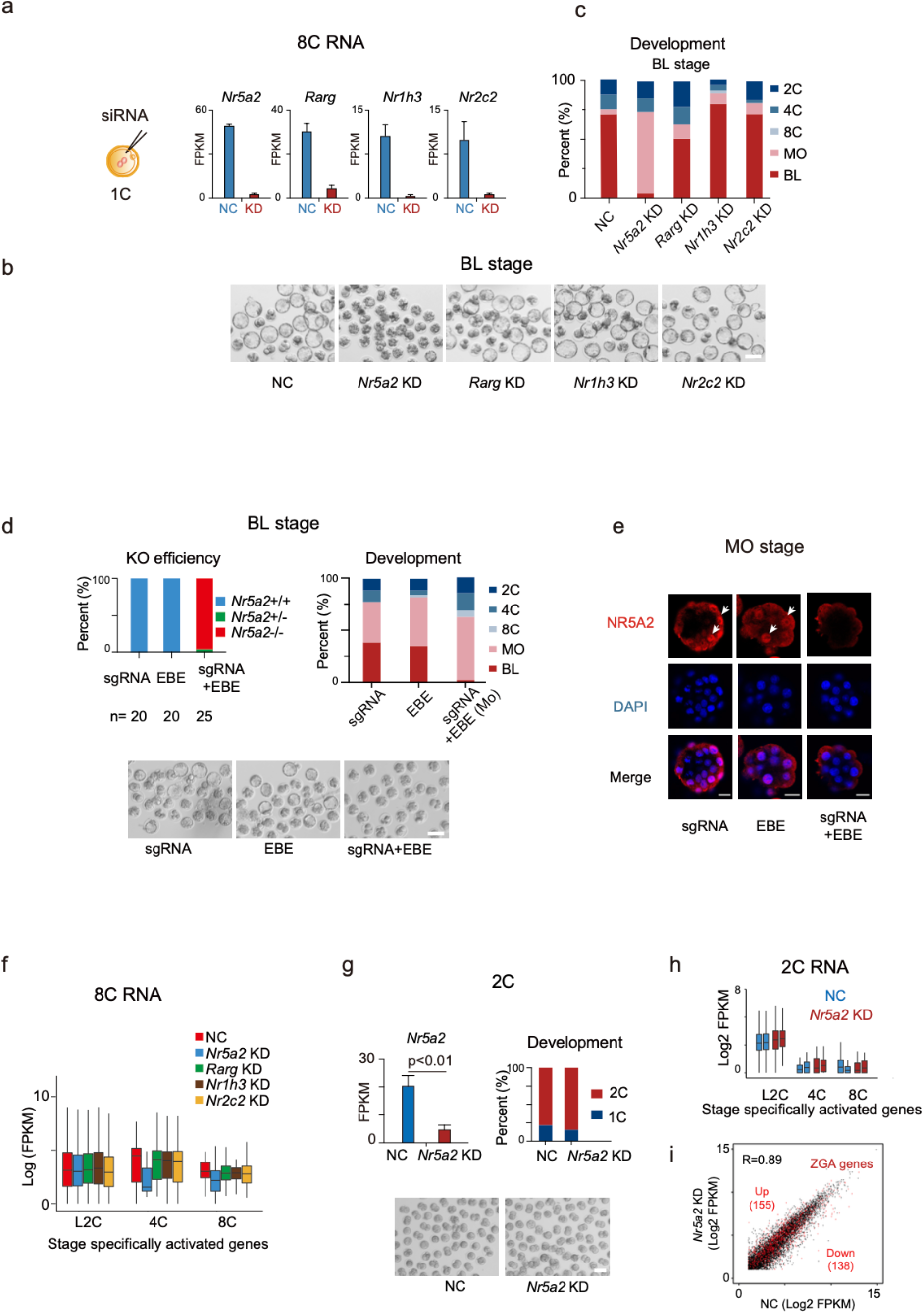
Both knockdown and knockout of *Nr5a2* led to morula arrest. **a**, Schematic of single knockdown (KD) of *Nr5a2, Rarg, Nr1h3*, and *Nr2c2* in the 1C embryos (left). Bar charts show the expression of *Nr5a2, Rarg, Nr1h3*, and *Nr2c2* in the negative control (NC) group (injected with negative control siRNA) and siRNA targeting group based on RNA-seq (right). The error bars denote the standard deviations of two biological replicates of RNA-seq. **b**, Embryo morphology after knocking down *Nr5a2, Rarg, Nr1h3*, or *Nr2c2* at the blastocyst stage. BL, blastocyst. Scale bar: 100 μm. **c**, Bar plots show the developmental rates of NC group, *Nr5a2* KD group, *Rarg* KD group, *Nr1h3* KD group, and *Nr2c2* KD group at the blastocyst stage (E4.5). MO, morula; BL, blastocyst. **d**, Bar charts show the percentages of base edited embryos after injection of *Nr5a2* sgRNA only, EBE mRNA only, and both sgRNA and EBE mRNA. Blue, unedited embryos. Green, one allele-edited embryos. Red, two allele-edited embryos (top, left). Bar plots show the developmental rates after injection of sgRNA only, EBE mRNA only, and both sgRNA and EBE mRNA at the blastocyst stage (E4.5) (top, right). Embryo morphology of different groups at the blastocyst stage (E4.5) are also shown (bottom). MO, morula; BL, blastocyst. **e**, Immunofluorescence of NR5A2 (red) and DAPI (blue) at the morula stage (E2.75) after injection of sgRNA only, EBE mRNA only, and both sgRNA and EBE mRNA. MO, morula. **f**, Box plots showing the average expression levels of stage specifically activated genes for NC groups (red), *Nr5a2* KD groups (blue), *Rarg* KD groups, *Nr1h3* KD groups, and *Nr2c2* KD groups (yellow) at the 8-cell stage. **g**, Bar charts showing the expression of *Nr5a2* in NC and *Nr5a2* KD group at the 2-cell stage (top, left), with *P* values (*t*-test, two-sided) indicated. Bar plots show the developmental rates of NC and *Nr5a2* KD groups at the 2-cell stage (top, right). Embryo morphology of NC and *Nr5a2* KD group at 2-cell stage is also shown (bottom). **h**, Box plots showing the average expression levels of stage specifically activated genes of NC groups (blue) and *Nr5a2* KD groups (red) at the 2-cell stage. The error bars denote the standard deviations of two biological replicates of RNA-seq. **i**, Scatterplot comparing gene expression of WT and *Nr5a2* KD in L2C embryos. ZGA genes are colored in red, and those that are upregulated and downregulated in *Nr5a2* KD embryos are indicated (2-fold change, adjust p-value <0.05).

### Genome-wide mapping of NR5A2 chromatin occupancy in mouse early embryos

To further investigate the direct targets of NR5A2 during mouse early development, we sought to capture its chromatin binding using CUT&RUN.^28^ We identified a sensitive NR5A2 antibody (Methods) that can detect its binding using as few as 1,000 WT mESCs, but not in *Nr5a2* knockout mESCs (Fig. S4a, b). NR5A2 motif was the top 1 enriched motif in its binding peaks (Fig. S4c), thus validating the data. Next, we performed NR5A2 CUT&RUN in the 2C and 8C embryos, which were highly reproducible (Fig. 3a, Fig. S4d). We could not detect NR5A2 binding in ICM likely due to its low expression (Fig. 1b) and the limited cells we collected. Therefore, we included mESCs for comparison instead. As a strong validation, the NR5A2 motif was again enriched in 2C and 8C binding peaks as the top 1 motif (Fig. 3b). At the 2-8C stages, NR5A2 showed distinct binding compared to that in mESCs and was preferentially enriched at the promoter of 2-8C specifically expressed genes (Fig. 3c; see Fig. 3d for example). In distal regions, NR5A2 also preferentially occupied sites specifically marked by H3K27ac and open chromatin,^13^ suggesting NR5A2 preferentially bound active regulatory regions (Fig. 3e). Stage-specific binding analysis showed that 2C specific and 2-8C shared NR5A2 distal binding were enriched near genes involved in housekeeping activities including DNA repair, RNA processing, histone modification, and DNA methylation (Fig. 3e, 1^st^ and 4^th^ clusters), suggesting activation of basic cellular machinery at the 2C stage. Interestingly, 8C-specific NR5A2 binding tended to enrich near genes functioning in blastocyst formation and stem cell maintenance, indicating the possible involvement of NR5A2 in lineage regulation at this stage (Fig. 3e, 2^nd^ cluster). By contrast, mESC-enriched binding was found near genes functioning in epithelial cell maintenance. LIF-responding genes also showed higher enrichment of NR5A2 in mESC (Fig. 3e, 3^rd^ and 5^th^ clusters). Notably, NR5A2 also bound the putative enhancers near *Nr5a2* itself (Fig. 3d), suggesting a positive-feedback regulatory loop. Taken together, these data demonstrate that we successfully captured NR5A2 binding in 2C and 8C embryos.

**Fig. 3.**
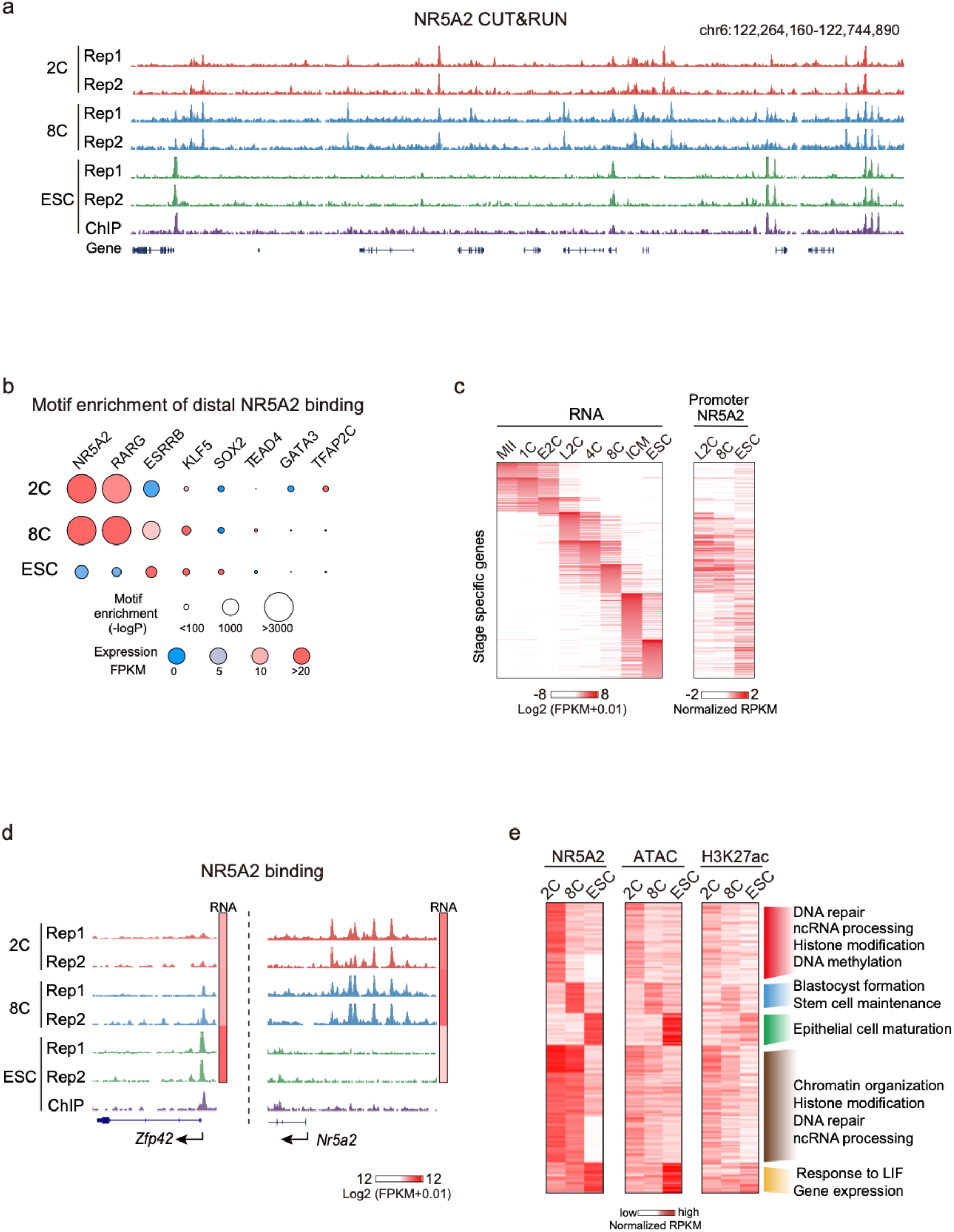
NR5A2 binding dynamics in mouse early embryos. **a**, The UCSC browser view showing NR5A2 CUT&RUN signals in 2-cell, 8-cell embryos, and mESCs (two biological replicates). NR5A2 ChIP-seq from a reference dataset ^20^ is also shown. **b**, TF motifs identified from NR5A2 distal binding peaks in 2-cell, 8-cell embryos, and mESCs. Sizes of circles indicate -Log P value. Expression levels of TFs are color coded. **c**, Heat maps showing stage-specific gene expression based on a reference dataset^13^ and their promoter NR5A2 binding enrichment. ICM, inner cell mass. **d**, The UCSC browser views and heat maps showing NR5A2 enrichment and gene expression of representative genes, respectively. **e**, Heat maps showing the NR5A2 binding, ATAC^13^, and H3K27ac signals at the stage-specific and shared NR5A2 binding peaks (left). The GO terms and example genes are also shown for different clusters (right).

### NR5A2 targets were preferentially down-regulated upon its deficiency

We then asked if NR5A2 regulates its binding targets by focusing on the 8C stage when both NR5A2 chromatin occupancy data and *Nr5a2* knockdown RNA-seq data were available. NR5A2 binding was preferentially enriched at the promoters of downregulated but not upregulated genes upon *Nr5a2* KD (Fig. 4a, left), suggesting that it functions primarily as an activator rather than a repressor. Consistently, distal NR5A2 binding peaks also preferentially resided near the down-regulated genes (Fig. 4a, right). Supporting a direct activation role of NR5A2 at these genes, those with NR5A2 binding and more NR5A2 motifs at promoters were more downregulated in *Nr5a2* KD embryos (Fig. 4b). Among 2,244 genes expressed at the 8C stage whose promoters were also occupied by NR5A2, a significant fraction (34%, n=263, p value<0.001) were downregulated (Fig. 4c). These data demonstrated that NR5A2 directly regulates gene activation at the 8C stage. For example, NR5A2 bound and activated *Eif4e* and *Cldn4* (Fig. 4d), which were involved in the transition of maternal to zygotic translation and tight junction regulation, respectively, and have been shown to be essential for peri-implantation development and normal blastocyst formation.^29, 30^ Moreover, it also bound pluripotency genes *Oct4, Nanog*, and *Tdgf1* which were all down-regulated in *Nr5a2* KD 8C embryos (Fig. 4d). Globally, we identified 2,108 genes that were specifically expressed in ICM compared to TE using a reference dataset ^13^, among them 74% (n= 1,557) were already expressed at the 8C stage (FPKM>1), which we termed early ICM genes (Fig. 4e, top). 187 of these early ICM genes were downregulated in *Nr5a2* KD 8C embryos, constituting 18% of all down-regulated genes (Fig. 4e, bottom). Importantly, the downregulated, but not the upregulated, early ICM genes were more enriched for NR5A2 binding at their promoters or in their neighbor distal regions (Fig. 5f, g), indicating they are part of direct targets of NR5A2. Together, these results suggest that NR5A2 directly promotes transcription at the 8C stage including a subset of pluripotency genes.

**Fig. 4.**
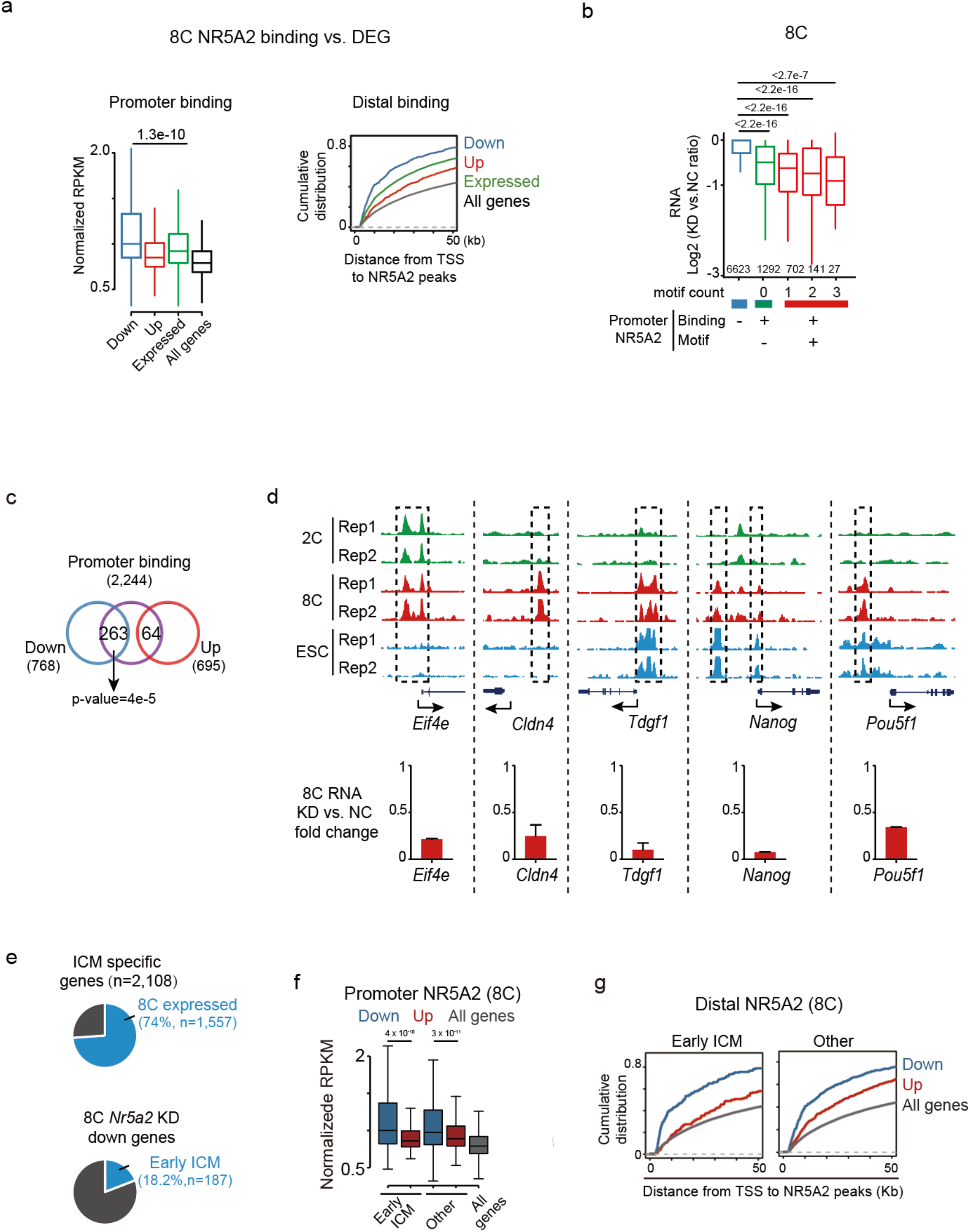
NR5A2 regulated the expression of its binding targets in early development. **a**, Boxplots showing the average enrichment of NR5A2 binding signals at the promoters (TSS ± 2.5 kb) (in WT embryos) of down-regulated, up-regulated, expressed, and all genes identified in *Nr5a2* KD 8C embryos (left), with P values (t-test, two-sided) indicated. The cumulative distributions of down-regulated, up-regulated, expressed, and all genes with defined distances (x-axis) between their TSSs and nearest distal 8-cell NR5A2 binding peaks are shown (right). **b**, Box plots showing the fold changes of gene expression in *Nr5a2* KD 8-cell embryos for all expressed genes based on the NR5A2 binding states and the numbers of motifs at promoters, with *P* values (*t*-test, two-sided) indicated. **c**, Venn diagram shows the overlap of down- and up-regulated genes in *Nr5a2* KD 8-cell embryos with NR5A2 target genes (with NR5A2 binding at promoters) (left). *P* values based on the hypergeometric distribution are shown. **d**, The UCSC browser views showing NR5A2 binding of representative genes in WT embryos and mESCs (top). Bar plots shows expression of representative genes in control and *Nr5a2* KD 8C embryos of two biological replicates of RNA-seq (bottom). **e**, Pie chart showing the percentages of ICM-specific genes that already show expression (FPKM>1) at the 8C stage (n=1,557) (top). Pie chart shows the percentages of down-regulated genes in *Nr5a2* knockdown 8C embryos that correspond to early ICM-specific genes (expressed at the 8C stage) (bottom). **f**, Boxplots showing the average enrichment of NR5A2 signals at the promoter (TSS ± 2.5 kb) (in WT embryos) for down-regulated or up-regulated genes identified in *Nr5a2* knockdown 8C embryos for ICM-specific and other 8C expressed genes, with *P* values (*t*-test, two-sided) indicated. All genes were similarly analyzed and are shown as controls. **g**, The cumulative distributions of down-regulated or up-regulated ICM-specific, TE-specific, and other genes in *Nr5a2* knockdown 8-cell embryos with defined distances (x-axis) between their TSSs and nearest distal 8-cell NR5A2 binding peaks.

**Fig. 5.**
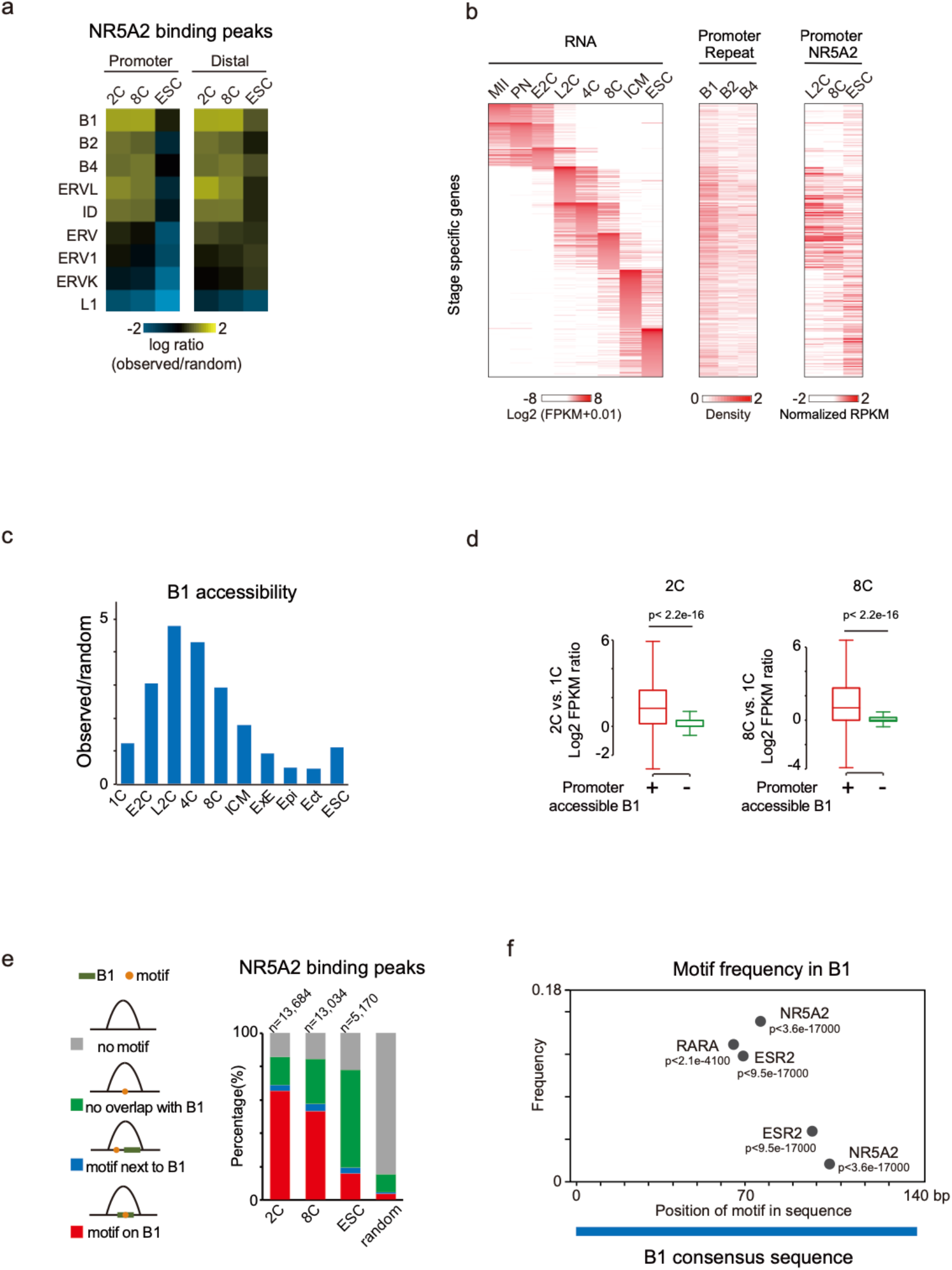
NR5A2 bound B1 repeats at the 2-8C stage. **a**, Heat maps showing enrichment (observed number/expected number from randomized peaks) of each repeat family in promoter and distal NR5A2 binding peaks. **b**, Heat maps showing the gene expression, promoter repeat density and NR5A2 binding enrichment at stage-specific genes. E2C, early 2-cell. L2C, late 2-cell. ICM, inner cell mass. **c**, Bar plots show the enrichment of accessible B1 repeats based on ATAC-seq and DHS-seq data relative to the background (observed number/expected number) in early embryos. E2C, early 2-cell. L2C, late 2-cell. ICM, inner cell mass; ExE, extraembryonic ectoderm; Epi, epiblast; Ect, ectoderm. **d**, Box plots showing expression activation levels (late 2C/1C (left) and 8C/1C (right)) for genes near accessible or inaccessible B1 at the 2C and 8C stage, with *P* values indicated. The error bars denote the standard deviations of two biological replicates of RNA-seq. **e**, Bar charts shows percentages of NR5A2 binding peaks at each stage in four types – motif in B1 (red), motif next to B1 (blue), motif non-overlapping with B1, and no motif (grey). **f**, TF motif frequencies and relative positions in the B1 consensus sequence are shown, with *P* values indicated.

### NR5A2 predominantly occupied the B1 repeats in the 2C and 8C embryos

Repetitive elements actively wire host transcriptional circuitry in part through functioning as cis-regulatory elements by providing transcription factor binding sites.^31^ Several classes of transposable elements are highly activated in mouse preimplantation embryos.^13, 32-35^ Interestingly, NR5A2 binding sites at the 2C and 8C stages, both for those at promoters and distal regions, overwhelmingly overlapped with the short-interspersed elements (SINEs) family, especially with the B1 repeats (Fig. 5a, Fig. S5a). For example, 84.8% and 73.3% of NR5A2 binding peaks at the 2C and 8C stages contained B1, respectively, compared to 18.9% of random peaks (Fig. S5a). B1 is abundantly expressed ^36^ and was indicated to play critical roles in mouse early embryos.^34^ B1 was preferentially present near 2-8C gene promoters, consistent with previous work,^37^ and correlated with NR5A2 binding at the 2C and 8C stages (Fig. 5b). B1 became widely accessible from the early 2C stage until the 8C stage based on ATAC-seq^13^ and DHS-seq^38^ data (Fig. 5c). Genes with accessible B1 at the promoters at the late 2C and 8C stages were more likely to be activated compared to inaccessible B1 (Fig. 5d). The accessible B1 at 2C stage was even more enriched for the NR5A2 motif (66.4%) compared to all B1 elements (52.5%). To further investigate the association of NR5A2 binding and B1 repeats, we classified the NR5A2 peaks into four groups based on the presence or absence of its motif and the B1 repeats, and whether the motif, if present, falls into B1 (Fig. 5e). At the 2C stage, the majority of NR5A2 binding peaks (64.9%) tended to contain B1 with the NR5A2 motif (Fig. 5e). This percentage then slightly decreased in 8C embryos (52.7%), and dropped abruptly in mESCs (15.6%). By contrast, NR5A2 preferentially bound its motif outside of B1 in mESCs (58.3% of all binding peaks) (Fig. 5e), suggesting the enriched binding of NR5A2 to B1 elements is specific for early embryos.

The data above raised two possibilities: the binding of NR5A2 may help open B1; alternatively, B1 may be pre-opened by other factors which facilitates the binding of NR5A2. The latter was supported by the fact that B1 became accessible starting from the early 2C stage (Fig. 5c), when *Nr5a2* expression was still low (Fig. 1b). Notably, the consensus sequence of B1 harbored motifs not only for NR5A2, but also for other TFs such as RARa and ESR (Fig. 5f). Similarly, other transposable elements such as B2 and B4 also possessed a set of TF motifs (Fig. S5b). Future studies are warranted to investigate if these TFs may bind and open these regions to allow the binding of NR5A2 to exert gene activation of downstream targets.

## Discussion

Transcription factors play essential roles in gene regulation and animal development. However, how TFs guide mammalian preimplantation development remains poorly understood. Here, through functional screening of TFs that were highly expressed after ZGA and showed motif enrichment in accessible chromatin, we identified NR5A2 as a key regulator of mouse early development and transcription program. The knockdown or knockout of *Nr5a2* led to preimplantation development defects with the 4-8C gene program impaired. Using CUT&RUN, we further determined possible direct targets of NR5A2 in early embryos, which included a panel of key pluripotency genes (i.e., *Nanog, Pou5f1*, and *Tdgf1*) at the 8C stage (Fig. 6). Intriguingly, NR5A2 bound extensively to the B1 elements at the 2-8C stages, but less so in mESCs. This coincides with the widespread opening of B1 in mouse embryos. These data demonstrate a critical role of NR5A2 in mouse early development by connecting ZGA to lineage segregation.

**Fig. 6.**
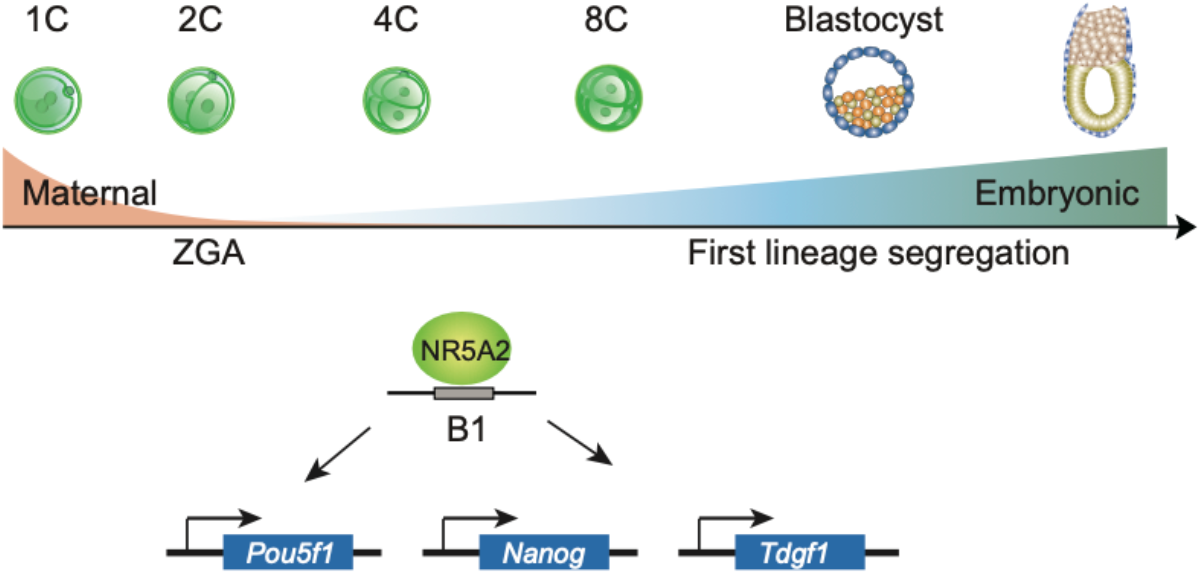
NR5A2 connects ZGA to the first lineage specification in early development. During mouse early development, NR5A2, a nuclear receptor TF highly induced during ZGA, predominantly binds B1 repeats and critically regulates 4-8C transcription (mid-preimplantation gene activation), including key lineage regulator genes, such as *Pou5f1, Nanog* and *Tdgf1*.

Despite a vast amount of knowledge on pluripotency core TFs and their targets, their upstream TF regulators in early embryos are still poorly characterized. The loss of *Nanog, Pou5f1*, or *Sox2* did not affect the formation of blastocysts in mice,^9, 10, 39^ suggesting that at least the initiation and commitment of the first lineage segregation does not strictly require these factors. Identification of upstream TFs is in part hindered by the limited research materials from early embryos and the lack of a cell line *in vitro* that can well recapitulate early embryo development, especially during the period of ZGA to morula. On the other hand, ATAC-seq and related methods coupled with RNA-seq provide a powerful tool to identify possible TF regulator candidates. As a testimony of this approach, NR5A2 represents one of the earliest TFs identified to date that exert functions on gene programs in early embryos starting as early as the 4C stage. Importantly, NR5A2 is required for the proper expression of a number of key pluripotency genes including *Pou5f1* and *Nanog* at the 8C stage. Interestingly, *Nr5a2* is downregulated in blastocyst (Fig. 1b) and it alone is dispensable for mESC self-renewal and pluripotency.^23, 24^ It is possible that when pluripotency core TFs are fully expressed, they form interconnected regulatory circuitry^8, 40^ and take over the pluripotency regulatory network, without requiring the continuous activation by NR5A2.

Of note, the *Nr5a2* KD and KO phenotypes in this study were more severe than those reported in previous studies.^21, 41^ Given different regions, with some corresponding to distinct isoforms of *Nr5a2*, were targeted in different KO strategies, it remained to be determined if the discrepancy may arise from these differences or other factors. Of note, a recent study reported a requirement of maternal NR5A2 in mouse ZGA and development beyond the 2C stage using a NR5A2 chemical inhibitor and *Nr5a2* knockdown in oocytes.^42^ By contrast, *Nr5a2* KD and KO starting from the 1C stage in our experiments did not have a major impact on ZGA and 2C development. While the expression of maternal *Nr5a2* is low (Fig. 1b), we could not assess its potential role in our study, which warrants future studies using the maternal and maternal-zygotic *Nr5a2* mutant mice.

Our NR5A2 CUT&RUN data in the 2C embryos also present one of the earliest TF regulomes studied thus far. These data provide strong functional support for previously identified putative regulatory elements marked by ATAC-seq and histone acetylation,^13^ as they interact with NR5A2 and the nearby genes were preferentially downregulated upon *Nr5a2* KD. Interestingly, NR5A2 pervasively occupied motifs embedded in accessible B1 elements in 2C and 8C embryos. Owing to the global epigenetic resetting after fertilization, the chromatin landscape of the mouse early embryo is extensively shaped by transiently expressed transposable elements, such as B1, B2, B4, and MERVL.^13^ Once regarded as the junk of the genome,^43, 44^ transposable elements are now considered as key players in gene regulation through cis- or trans-regulatory mechanisms.^31^ These functions are in part mediated by their embedded TF motifs.^45, 46^ For example, the B2 repeats possess the CTCF motif that can actively wire chromatin organization.^47, 48^ In mouse embryos, the knockdown of B1 induced 2C arrest,^49^ although the underlying mechanisms remain unclear. Given that B1 elements are already accessible prior to major ZGA^3^ (Fig. 5c) and NR5A2 is strongly induced after ZGA (late 2C stage) (Fig. 1b), we favor the possibility that B1 may be opened by other TFs at least at the 2C stage, which further allows the binding of NR5A2 that subsequently triggers downstream events including the activation of pluripotency genes. This may also explain why *Nr5a2* KD has a limited impact on ZGA. We nevertheless cannot rule out the possibility that NR5A2, once expressed, may in turn, participate in future opening these elements. B1 activation likely requires multiple factors, as B1 elements are rarely bound by NR5A2 and are mostly inaccessible in mESCs. Given that it is technically challenging to perturb B1 in early embryos, the relationship between B1 activities and the binding of NR5A2 as well as other TFs remains to be further explored. In sum, we identified NR5A2 as a critical TF regulator for the 4-8C stage-specific transcription in mouse embryos, thus filling a major gap of transcription circuitry in early development that connects ZGA to lineage segregation. The TF-regulome maps presented here also pave the ways for future studies to decode transcription circuitry underlying early development and cell fate decisions when life begins. Finally, given *Nr5a2* expression is quite low in human embryos unlike its counterpart in mouse ^50, 51^ (Fig. S6a), it would be highly interesting to investigate whether other nuclear receptor TFs may execute similar functions in human early development.

## Acknowledgments

We are grateful to members of the Xie laboratory for discussion and comments during the NR5A2 study and the preparation of the manuscript, and the Animal Center and Biocomputing Facility at Tsinghua University for their support. We are grateful to Drs. Pablo Navarro Gil, Nicola Festuccia, and Michel Cohen-Tannoudji for sharing unpublished data and insightful discussion. We are grateful to Dr. Hui Yang for sharing the reagents and Dr. Wanlu Liu for discussion.

## Author contributions

F.L., L.Li, and W.X. conceived and designed the project. F.L. performed embryo experiments with the help from L.Liu. F.L. performed microinjection and immunostaining with the help of L.Liu. L.Li performed CUT&RUN. X.H. and Z.Z. performed RNA-seq. X.H. constructed the Nr5a2 KO mESC. F.L. and L.Li performed the bioinformatics analysis. F.L., L.Li., B.L., X.H., and W.X. prepared most figures and wrote the manuscript with help from all authors.

## Supplementary Materials for

### METHODS

#### Data reporting

No statistical methods were used to the predetermine sample size. The experiments were not randomized and the investigators were not blinded to allocation during outcome assessment.

#### Animal maintenance

WT C57BL/6J strain mice were purchased from Vital River. PWK/PhJ mice were originally purchased from Jackson Laboratory and raised at Tsinghua Animal Center. Mice were maintained under SPF conditions with a 12 h-12 h light-dark cycle in a 20-22°C environment. All animals were taken care of according to the guidelines of the Institutional Animal Care and Use Committee (IACUC) of Tsinghua University, Beijing, China.

#### Early embryo collection

To obtain mouse preimplantation embryos, 5-6-week-old C57BL/6N female mice were intraperitoneally injected with pregnant mare’s serum gonadotropin (PMSG; 5 IU) and human chorionic gonadotrophin (hCG; 5 IU) 44-48 hours after PMSG injection. After mating with PWK/PhJ males (Jackson Laboratory), preimplantation embryos were collected in M2 medium at the following time points after hCG injection: 43 h (late 2-cell) and 68-70 h (8-cell).

#### Cell culture and *Nr5a2* knockout mESC generation

Mouse ES cells (mESC) were cultured on 0.1% gelatin pre-coated dish in DMEM (Gibco Cat. 11995-065) containing 15% FBS (Hyclone Cat. SH30396.03), leukemia inhibiting factor (LIF) (Millipore Cat. GSE1107), penicillin/streptomycin (Millipore Cat. TMS-AB2-C), GlutaMAX (Gibco Cat. 35050-061), nucleosides (Millipore Cat. ES-008-D), β-mercaptoethanol (Gibco Cat. 21985-023), and non-essential amino acids (Gibco Cat. 25-025-CIR). To target the *Nr5a2* gene, four sgRNA sequences (ctaagaatgtctgctagtt, agccactctcttaacatc, aaggcctgaacttgtgta, aaggcctgaacttgtgta) were cloned into a pX330 plasmid (Addgene, 42230) and were co-transfected into the R1 mESCs with Lipofectamine 3000 (Thermo Fisher Scientific). Two to three days after transfection, cells were manually sorted into a gelatinized 96-well plate for single-clone selection. The obtained clones were genotyped by PCR and validated by Sanger sequencing. The homozygous *Nr5a2* knockout cell lines were further validated by western blot for NR5A2.

#### Immunostaining

All steps were performed at room temperature (RT). Mouse embryos were fixed with 4% paraformaldehyde (PFA) (Sigma-Aldrich, P6148) for 30 mins and then permeabilized with 0.5% Triton X-100 in PBS for 30 min. The samples were blocked with 1% BSA for 1 h and incubated with primary antibodies (1:200) for 1 hour. The primary antibody was washed out with PBST (0.1% Triton X-100 in PBS) and then incubated with the secondary antibody and Hoechst 33342 for 30 min. The samples were washed with PBST three times. All immunofluorescence images were taken by LSM880 confocal microscope system (Zeiss) and were analyzed using ImageJ.

#### *In vitro* transcription and base editor

The sgRNA (ggtccgatcgcatggggaac) was cloned into pX330 vectors (Addgene, 42230). The primers ttaatacgactcactatagggtccgatcgcatggggaac and aaaagcaccggactcggtgcc were used to obtain linearized sgRNA with T7 promoter. The pCMV-hA3A-eBE-Y130F (Addgene #113423) plasmids were linearized. The linearized sgRNA and eBE plasmids were then transcribed with T7 mMESSAGE Kit (Invitrogen, AM1334) following the manufacturer’s instructions. mRNAs were recovered by RNAClean XP beads (Beckman, A63987). 100 ng/μl for sgRNA and eBE mRNA were used. All injections were performed with an Eppendorf Transferman NK2 micromanipulator.

#### Nuclear receptor TF knockdown

To knock down four NR family members *Nr5a2, Rarg, Nr1h3*, and *Nr2c2*, three siRNAs were designed for each gene. The siRNA of negative control is UAAGGCUAUGAAGAGAUACTT. The siRNAs of *Nr5a2* are CCUCUGCAAUUCAGAACAUTT, GCUCACCUGAGUCAAUGAUTT, GGAGUGAGCUCUUGAUUCUTT. The siRNAs of *Rarg* are GCCAUGCUUUGUAUGCAAUTT, GGAAGCUGUAAGGAACGAUTT, CUUGUCUGGACAUCCUAAUTT. The siRNAs of *Nr1h3* are CCAUUCAGAGCAAGUGUUUTT, GGCUGCAACACACAUAUGUTT, GUGCAGGAGAUUGUUGACUTT. The siRNAs of *Nr2c2* are GCUCAUGAGCUCCAACAUATT, GCCAGAGUACCUCAAUGUATT, GCAAAUGUAGUGACCUCUUTT. For microinjection in each knockdown group, 1.67μM of each siRNA was used. For the negative control group, 20μM of siRNA was used. siRNAs were ordered from GenePharma.

#### RNA-seq library preparation and sequencing

All RNA-seq libraries were generated following the Smart-seq2 protocol as described previously.^1^ The zona pellucida was gently removed by treatment with Tyrode’s solution (Sigma, T1788). 5-10 embryos per sample were washed three times in M2 medium and then lysed in 2 μl lysis buffer containing RNase inhibitor.

#### CUT&RUN library generation and sequencing

CUT&RUN was conducted following the published protocol.^2^ ESCs or embryos with zona pellucida removed were transferred into a 0.2mL conventional, non-low-binding PCR tube (Axygen). The samples were then resuspended by 60μL washing buffer (HEPES-KOH, pH = 7.5, 20mM; NaCl, 150mM; Spermidine, 0.5mM, and with Roche complete protease inhibitor). 10-20μL Concanavalin-coated magnetic beads (Polyscience, 86057) for each sample were gently washed, resuspended by binding buffer (HEPES-KOH, pH = 7.9, 20mM; KCl, 10mM; CaCl_2_, 1mM; MnCl_2_, 1mM) and carefully added to the samples. The samples were then incubated at 23°C for 10 mins on Thermomixer (Eppendorf) at 400 rpm. The samples were then held at the magnetic stand to carefully aspirate buffer and were resuspended by 60μL antibody buffer (washing buffer plus 0.01% digitonin and EDTA, pH = 8.0, 2mM) with NR5A2 antibody diluted at a ratio of 1:50. The samples were then incubated at 4°C on Thermomixer for 3 hours at 400 rpm. After that, the samples were held at the magnetic stand and then resuspended by the secondary antibody to a final concentration of 1:100 at 4°C on Thermomixer for 1 hour at 400 rpm. The samples then were placed the tube on the magnet stand to remove the liquid and added 75μL dig washing buffer with pA-MNase (to a final concentration of 700ng/mL), and incubated at 4°C on Thermomixer for 1 h at 400 rpm. The samples were washed with 200μL dig washing buffer twice at the magnetic stand. The samples were resuspended by 100μL dig washing buffer and balanced on ice for 2 mins. Targeted digestion was performed on ice by adding 2μL 100mM CaCl_2_ for 30 mins, and reaction was stopped by adding 100μL 2 × stop buffer (NaCl, 340mM; EDTA, pH = 8.0, 20mM; EGTA, pH = 8.0, 4mM; RNase A, 50μg/mL; glycogen, 100μg/mL) and fully vortexed. The samples were then incubated at 37°C for 30-45 min for fragment release. The supernatants were purified by phenol-chloroform followed by ethanol purification.

Purified DNA was subjected to Tru-seq library construction using NEBNext Ultra II DNA Library Prep Kit for Illumina (NEB, E7645S). The DNA was resuspended in 25μL ddH2O, followed by end-repair/A-tailing with 3.5μL End Prep buffer and 1.5μL enzyme mix according to manufactory instructions. The ligation reaction was then performed by adding diluted 1.25μL 0.25μM adaptors (NEB Multiplex Oligos for Illumina, E7335S), 15μL Ligation master mix, and 0.5μL Ligation enhancer, at 4°C for overnight or 20°C for 2 h. The ligation reaction was treated with 1.5μL USERTM enzyme according to the instruction and was purified by 1.4 × AMPure beads. The PCR was performed by adding 25μL 2 × KAPA HiFi HotStart Ready Mix (KAPA Biosystems, KM2602) with primers of NEB Oligos kit, with the program of 98°C for 45s, (98°C for 15s and 60°C for 10s) with 16 cycles and 72°C for 1 min. The final libraries were purified by 1.4 × AMPure beads and subjected to next-generation.

#### Western blot

The samples were lysed with 2X Tris-glycine-SDS sample buffer including 1X Roche complete protease inhibitor. Samples were heated at 99 °C for 5 min and loaded on a BioRad 4-15% gradient Tris-glycine gel, then transferred to low fluorescence PDVF membrane using a BioRad System. Blots were briefly rinsed in TBS, blocked in 5% non-fat milk in TBST for 1 h, and incubated overnight at 4 °C in primary antibody (1:1,000). Blots were washed in TBST, incubated for 1 h at room temperature in peroxidase-conjugated anti-rabbit or anti-mouse IgG (Jackson ImmunoResearch) diluted 1:10,000 in 1% non-fat milk in TBST, then washed in TBST. The signal was detected with SuperSignalTM West Femto Maximum Sensitivity Substrate (Thermo Scientific) and imaged using a BioRad ChemiDocTM Touch Imaging system.

#### Data analyses

##### RNA-seq data processing

Paired-end RNA-seq reads were trimmed and mapped to the mm9 genome by TopHat v2.1.1. ^3^ Cufflinks v2.2.1 ^3^ was used to calculate the FPKM per gene based on mm9 refFlat from the UCSC genome annotation database^4^. Differential expression analysis was performed with R package edgeR. Genes with FDR <0.01 and log2(fold change) >=2 were considered as differentially expressed.

#### Identification of stage-specific genes and ZGA genes

ZGA genes and stage-specific genes were defined based on the reference RNA-seq data using staged mouse embryos dissected in *vivo*. ZGA genes were defined as those not expressed or lowly expressed in FGO and MII oocytes (FPKM < 5) but become upregulated (FPKM > 5, at least 3-fold upregulation) in either early 2-cell embryos or late 2-cell embryos. Genes that are expressed in oocytes (FPKM > 5) but are highly upregulated at the late 2-cell stage (over 5-fold upregulation) were also included in major ZGA genes (n=99). No such genes exist for minor ZGA genes. Maternal genes were defined as those that are expressed in MII or FGO oocytes (FPKM > 5) but become downregulated (at least 3-fold) at the late 2-cell stages. Genes specifically activated at each stage during early development were defined using more strict criteria to ensure their stage specificity. These genes are activated at a defined stage (FPKM > 5) but stay silenced at all preceding stages from FGO (FPKM < 1).

#### CUT&RUN, ChIP-seq, and ATAC-seq data processing and peak calling

The paired-end reads were aligned with the parameters: -t –q –N 1 –L 25 –X 1000 --no- mixed --no-discordant by Bowtie2 (version 2.2.2).^5^ All unmapped reads, non-uniquely mapped reads, and PCR duplicates were removed. For downstream analysis, we normalized the read counts by computing the numbers of reads per kilobase of bin per million of reads sequenced (RPKM) for 100-bp bins of the genome. To minimize the batch and cell type variation, RPKM values across the whole genome were further Z-score normalized. To visualize the CUT&RUN signals in the UCSC genome browser, we generated the RPKM values on a 100bp-window base. Peaks were called using MACS v1.4.2 ^6^ with the parameters nolambda –nomodel. Promoters were defined as ±2.5 kb around the transcription starting sites (TSS). Peaks at least 2.5 kb away from TSSs were defined as distal peaks by BEDTools v2.29.0.^7^

#### Binding site feature annotation and motif analyses

The findMotifsGenome.pl script from HOMER program^8^ was used to identify the enriched motifs at enhancers. AnnotatePeaks.pl was then used to identify specific peaks that contain certain motifs. Motif densities at 10 bp resolution within 500 bp around peak center were analyzed using the annotatePeaks.pl with the parameters: annotatePeaks.pl mm9 - size -500,500 -hist 10.

#### GO analysis

The DAVID web-tool was employed to identify the GO terms using databases including Molecular Functions, Biological Functions and Cellular Components.^9^ The functional enrichment for genes that are near stage-specific distal peaks was analyzed using the GREAT tool by default settings.^10^

#### Identification of stage-specific NR5A2 peaks

In brief, distal peaks from all stages were pooled, and average Z-score normalized RPKM scores were calculated in these regions. MAnorm2 ^11^ was used to identify differential peaks in any combinations of pairs of all examined stages. Peaks were further selected by the following criteria: the distal peak had high enrichment at this stage (normalized RPKM > 1) and positive enrichment (normalized RPKM > 0) was observed at no more than two additional stages.

#### The comparison between NR5A2 peaks and repetitive elements

To identify the enrichment of repetitive elements in promoter or distal NR5A2 peaks, the NR5A2 peaks were compared with the locations of annotated repeats (RepeatMasker) downloaded from the UCSC genome browser. As repeats of different classes vary greatly in numbers, a random set of peaks with identical lengths of CUT&RUN peaks were used for the same analysis as a control. The number of observed peaks that overlap with repeats were compared to the number of random peaks that overlap with repeats, and a log ratio value (log2) was generated as the “observed/expected” enrichment.

#### Enhancer vs. gene expression analysis

To identify the potential targeted genes for each putative enhancer (distal peaks), the nearest distal peaks from the TSS of the gene (within 50Kb) were assigned to this gene as its putative enhancers.

**Fig. S1.**
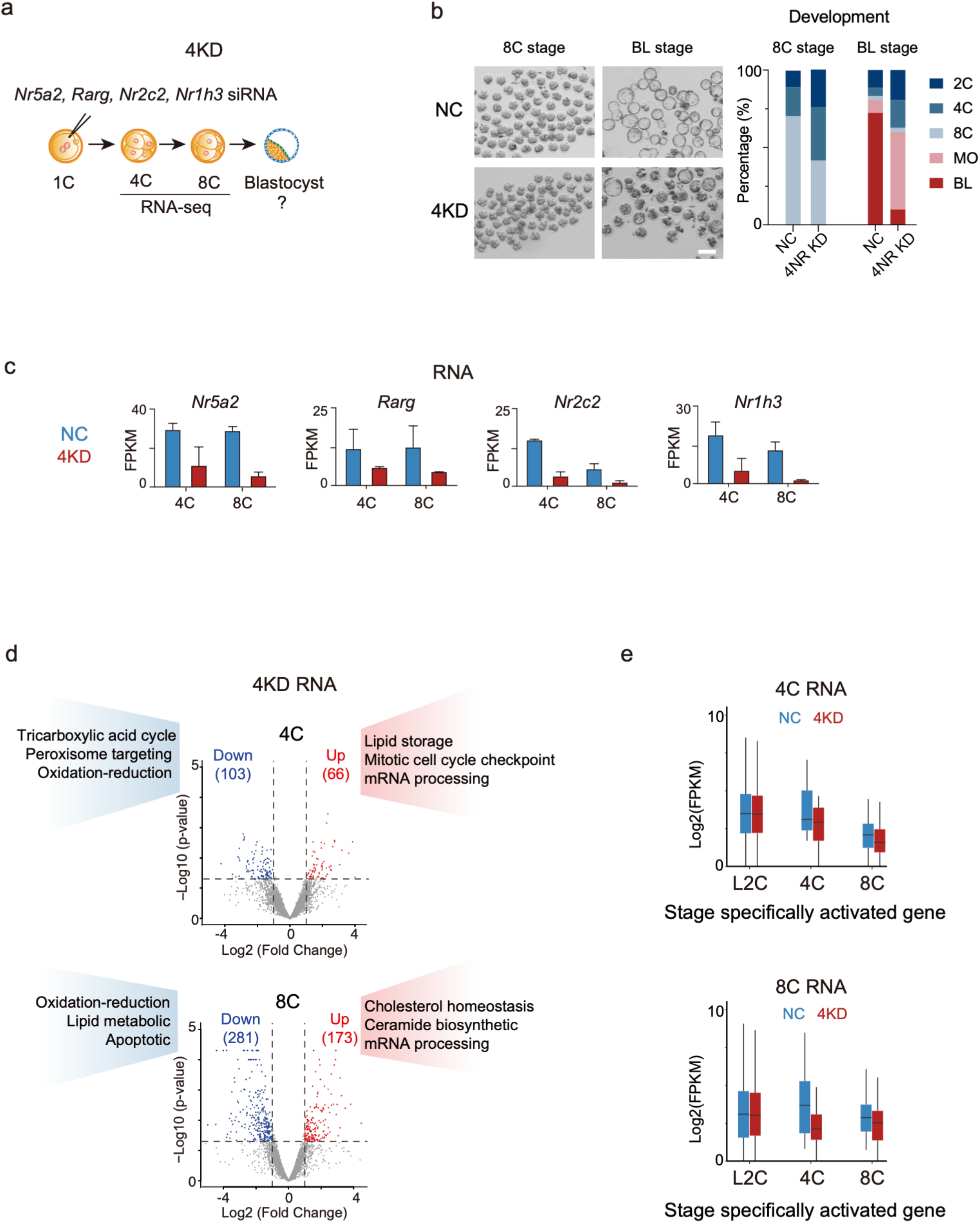
Joint knockdown of nuclear receptor TFs led to morula arrest. **a**, Schematic of the combined knockdown of 4 ZGA-activated nuclear receptor factors (4KD, knockdown of *Nr5a2, Rarg, Nr1h3*, and *Nr2c2*). **b**, Embryo morphology after knocking down 4 NR factors at the 8C and blastocyst stage (E4.5) (left). Scale bar: 100 μm. Bar plots show the developmental rates of the NC group and 4KD group at the blastocyst stage (E4.5) (right). MO, morula; BL, blastocyst. **c**, Bar charts showing the RNA expression of 4 nuclear receptor factors in NC and 4KD group based on RNA-seq. The error bars denote the standard deviations of two biological replicates of RNA-seq. **d**, Volcano plots showing differentially expressed genes in the 4C (top) and 8C (bottom) 4KD embryos. Up-regulated (log2(Fold change)>2, P value<0.05) and down-regulated (log2(Fold change)>2, P value<0.05) genes are colored in red and blue, respectively. The GO terms are also shown. **e**, Boxplots showing the average RNA expression levels of stage specifically activated genes in NC and 4KD groups at the 4C and 8C stages.

**Fig. S2.**
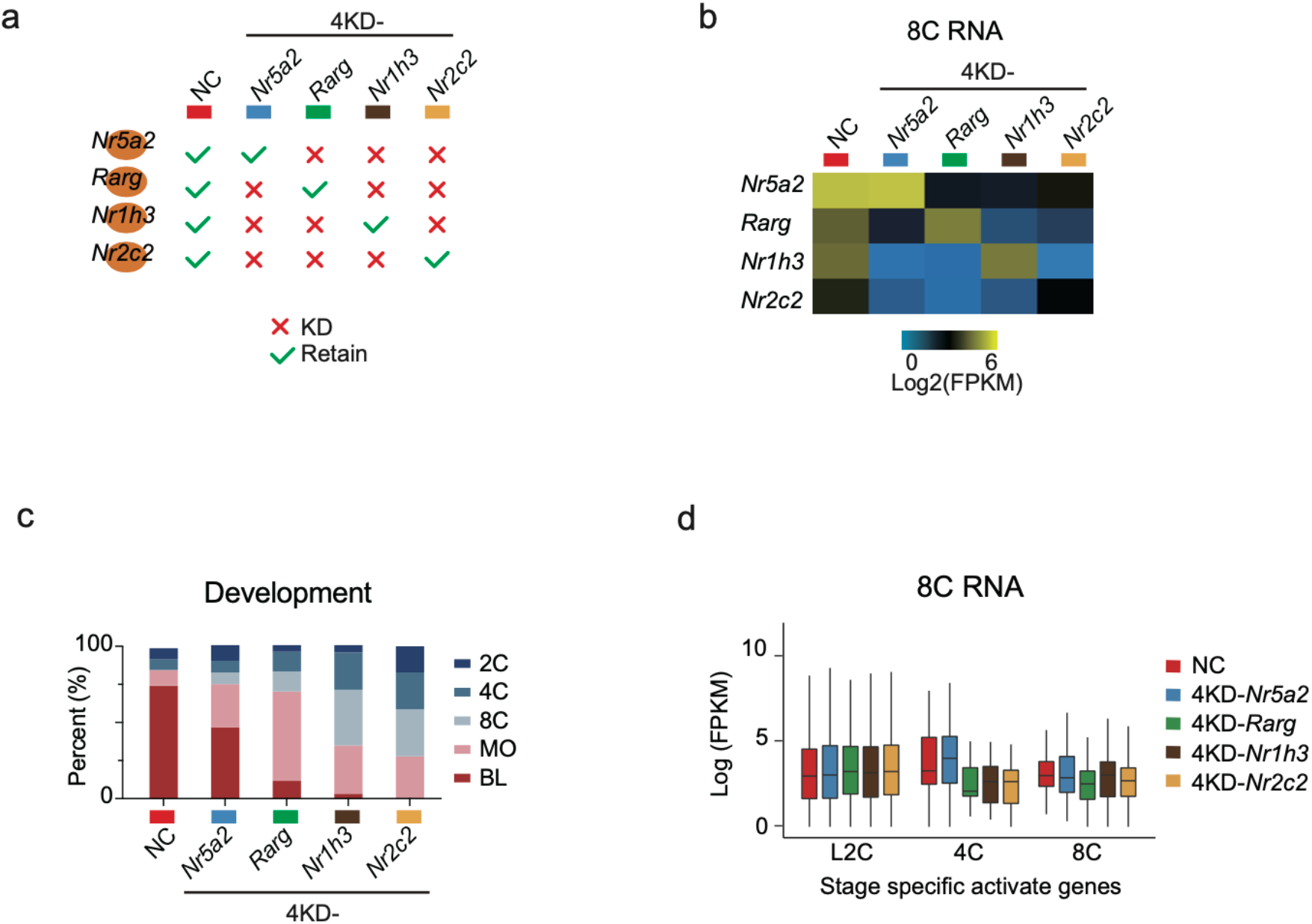
NR5A2 is the major functional factor among the 4 NR TFs activated during ZGA. **a**, Schematic illustration of the triple-knockdown (3KD, or 4 KD leave-one-out, 4KD-) among the 4 NR factors (*Nr5a2, Rarg, Nr1h3*, and *Nr2c2*). **b**, Heat maps showing expression levels of each NR factor in NC group and each 3KD group at the 8C stage. **c**. Bar charts showing the developmental rates of NC and the 3KD groups at the blastocyst stage (E4.5). **d**, Box plots showing the average RNA expression levels of stage specifically activated genes of NC group (red) and each 3KD group at the 8C stage.

**Fig. S3.**
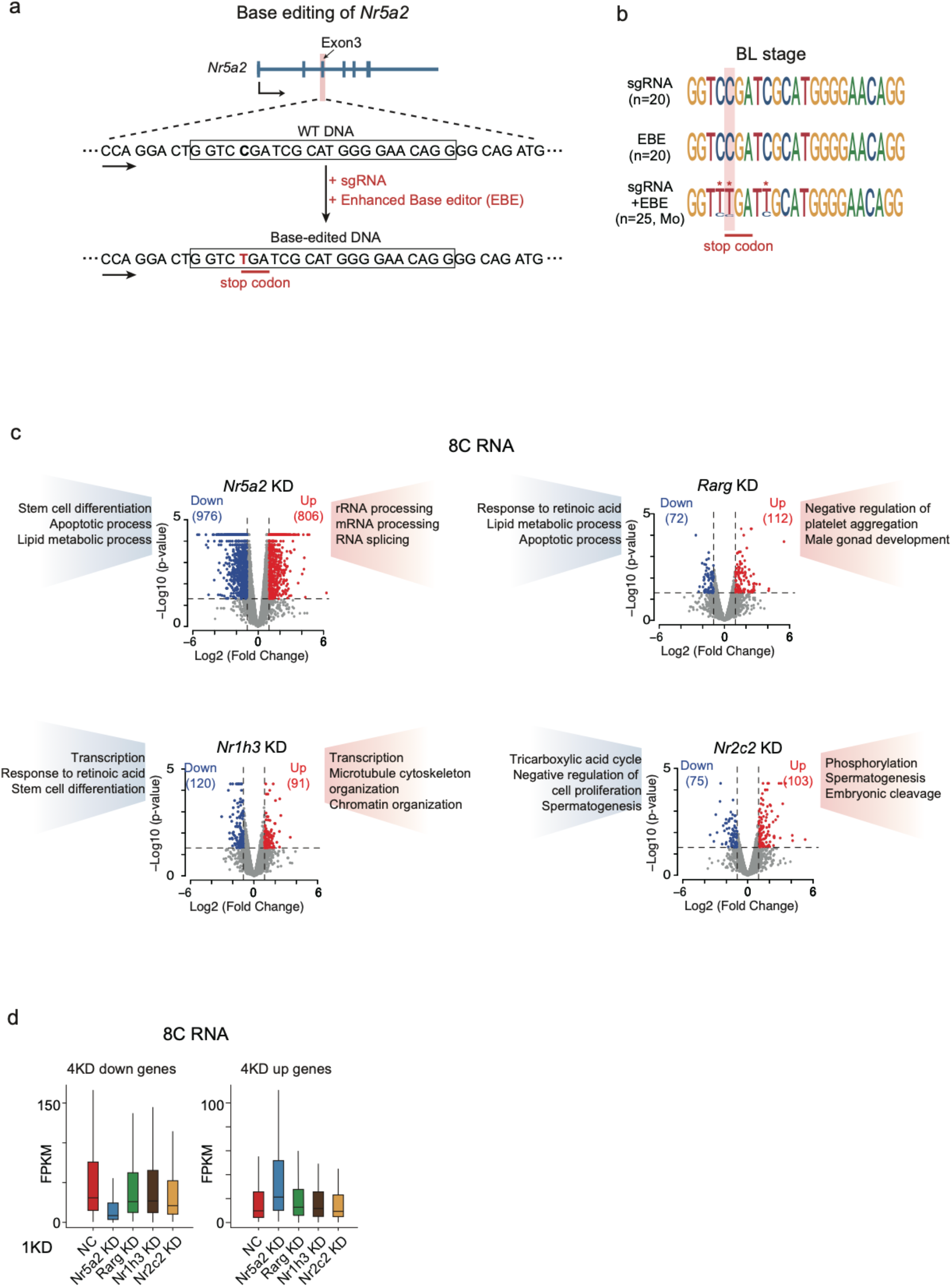
*Nr5a2* KO in mouse embryos. **a**, Schematic of base editing for *Nr5a2*. Exon 3 of *Nr5a2* was targeted (top). Box shows sgRNA targeting sequence (middle). Introduced stop codon is indicated (bottom). **b**, Sequence logos showing single-embryo genotyping results by Sanger sequencing for the sgRNA targeting region after injection of sgRNA only, EBE mRNA only, and both sgRNA and EBE mRNA. Asterisks denote the sites mutated by base editing, including two neighbor sites besides the stop codon. **c**, Volcano plots showing differentially expressed genes in *Nr5a2* (top, left), *Rarg* (top, right), *Nr1h3* (bottom, left), or *Nr2c2* (bottom, right) single KD embryos at the 8C stage. GO analysis results of differentially expressed genes are also shown. **d**, Box plots showing the average RNA expression levels for 4KD down-regulated and up-regulated gene lists in the NC group (red), *Nr5a2* (blue), *Rarg* (green), *Nr1h3* (brown), and *Nr2c2* single KD group (yellow) at the 8C stage.

**Fig. S4.**
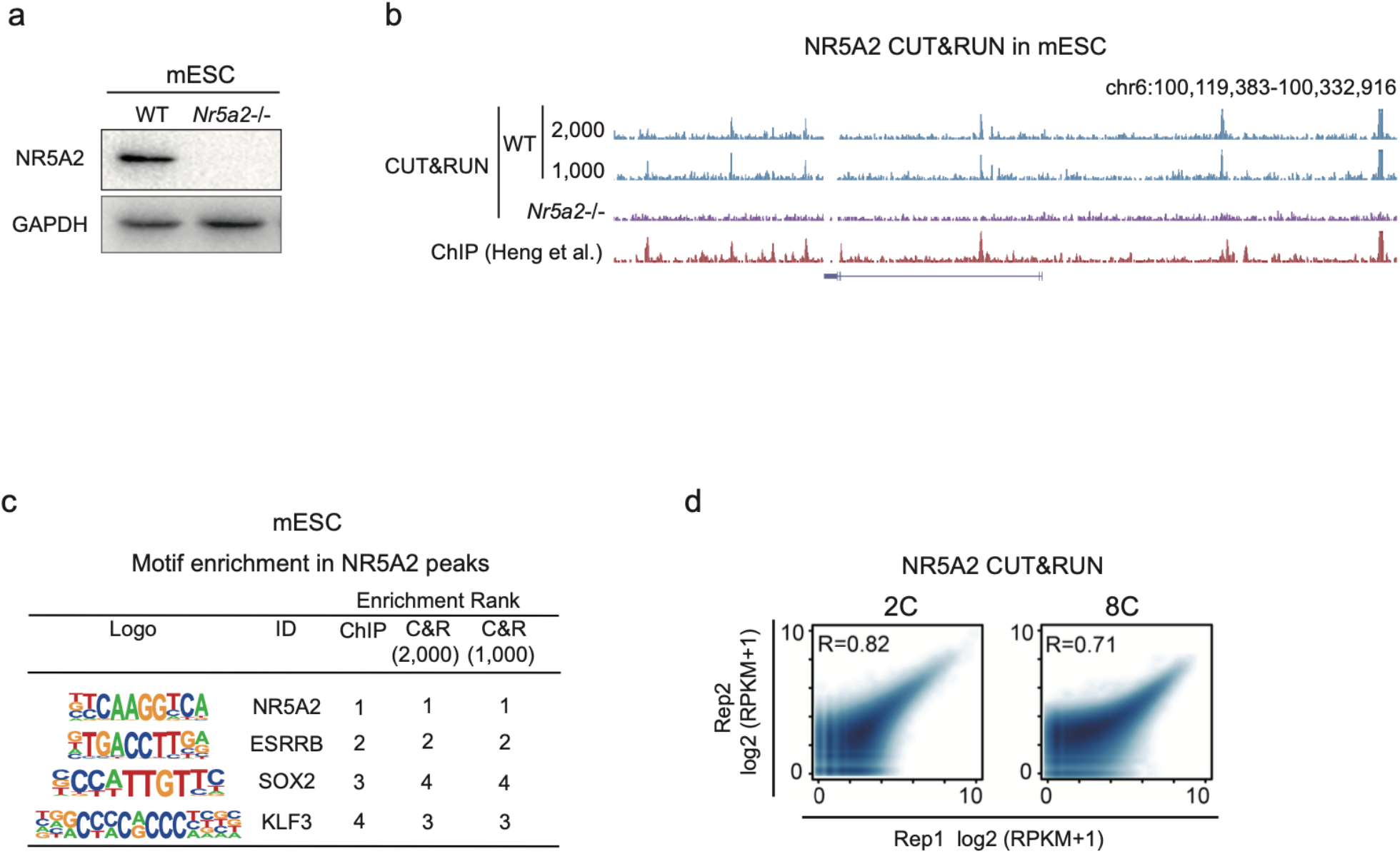
Validation of the NR5A2 CUT&RUN data. **a**, Western blot showing NR5A2 and GAPDH expression in WT and *Nr5a2*^*-/-*^ mESCs. **b**, The UCSC browser view showing NR5A2 CUT&RUN and ChIP-seq ^12^ signals in mESCs with cell numbers ranging from 2,000 to 1,000. **c**, Sequence logos and enrichment ranks of the motifs enriched in NR5A2 CUT&RUN peaks in mESCs. **d**, Scatter plots comparing the replicates of NR5A2 CUT&RUN data in 2C and 8C embryos. The Pearson correlation coefficients are also shown.

**Fig. S5.**
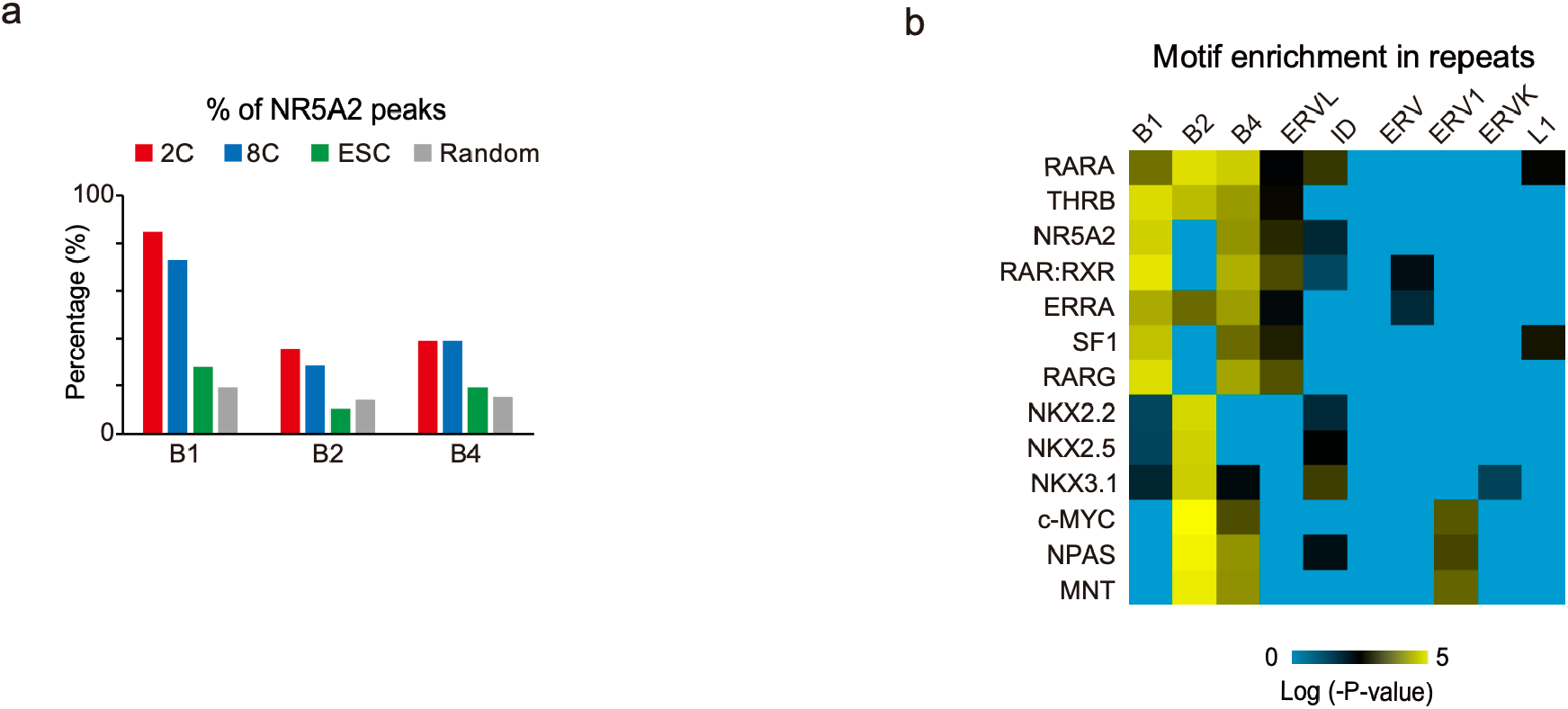
NR5A2 binding sites in early embryos enriched for transposable elements. **a**, Bar charts showing the percentages of NR5A2 peaks overlapped with B1, B2, and B4 repeats in 2C (red), 8C (blue), and ESC (green). Random peaks were shuffled and generated with lengths matched, and were similarly analyzed. **b**, Heat maps showing TF motif enrichment in different groups of repeats.

**Fig. S6.**
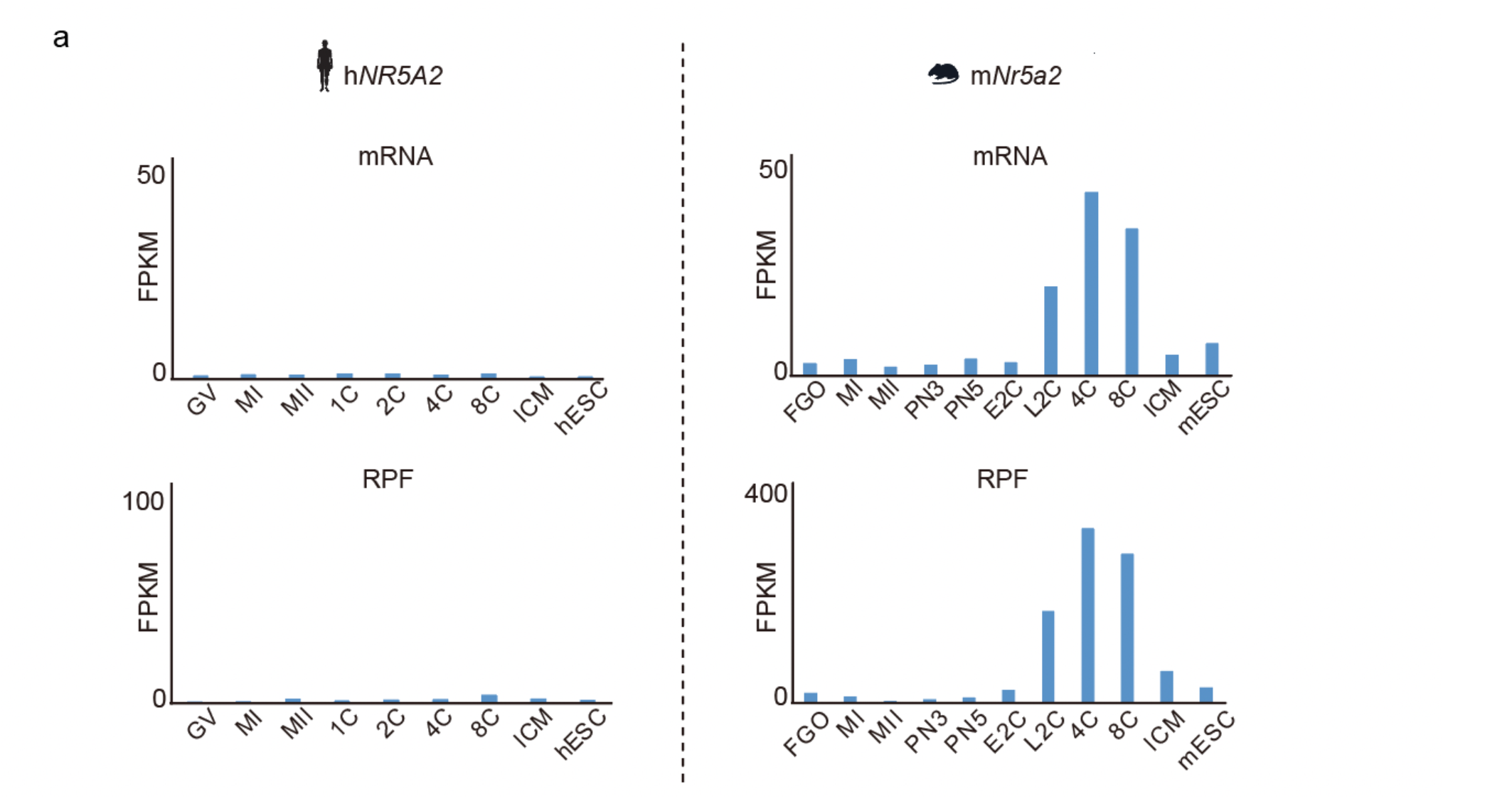
*NR5A2* expression in human and mouse early embryos. **a**, Bar charts showing *NR5A2/Nr5a2* mRNA levels from RNA-seq and ribosome-protected fragment (RPF, reflecting translation) levels from Ribo-seq in human (left) and mouse (right) early embryos.^13, 14^

